# Development of a polymicrobial checkerboard assay as a tool for determining combinatorial antibiotic effectiveness in polymicrobial communities

**DOI:** 10.1101/2022.10.12.511785

**Authors:** Caroline Black, Hafij Al Mahmud, Sabrina Wilson, Allie C. Smith, Catherine A. Wakeman

## Abstract

**AIMS:** To establish a methodology for identifying the effects of combinatorial antibiotic treatment on individual members of a polymicrobial community.

**METHODS AND RESULTS:** Both gentamicin and ceftazidime were diluted to concentrations ranging from 0.06 μg ml^-1^ to 128 μg ml^-1^. An equal ratio polymicrobial community of *Staphylococcus aureus, Pseudomonas aeruginosa, Enterococcus faecalis*, and *Acinetobacter baumannii* was inoculated into the combined antibiotics in the checkerboard and incubated for 18 hours in static conditions. After incubation, visible turbidity of the overall community was recorded, and bacteria from the wells were diluted to 10^-3^ and then spot plated on selective and differential media. After 24 hours, colony-forming unit (CFU) counts were obtained for each species.

**CONCLUSIONS:** Visible turbidity is not truly indicative of cell viability, and the polymicrobial community can decrease the antibiotic susceptibility of *P. aeruginosa*, rendering the clinically-established beneficial combination of gentamicin and ceftazidime ineffective.

**SIGNIFICANCE:** Previous checkerboard methodology which focuses on using visible turbidity to determine monomicrobial antibiotic susceptibility fails to account for polymicrobial cooperation that has been shown to reduce antibiotic efficacy. Our new methodology could be implemented in clinical microbiology laboratories with minimal impact on the overall time for diagnosis.

## INTRODUCTION

Antibiotic resistance is one of the biggest challenges faced when attempting to effectively treat patient infections. In certain cases, the initial treatment prescribed can mean the difference between a patient living or dying (Micek *et al*. 2005). In some cases, antibiotic combination therapy, in which two or more antibiotics are prescribed, has been shown to help clear infections quickly and more effectively than standard mono-treatment (Angst *et al*. 2021). Antibiotic combination therapy can be used if the infection requires a broader spectrum of therapy, if the infection is determined to be polymicrobial in nature, if antibiotic synergism is deemed helpful for treating the infection, or if the emergence of antibiotic resistance is seen while treating the infection (Ahmed *et al*. 2014). Currently, the decision regarding which antibiotics to use when treating an infection is based on empiric and definitive antimicrobial therapy (Leekha *et al*. 2011). Empiric antimicrobial therapy is when an antibiotic (or antibiotics) is chosen by a clinician based on the presentation of symptoms when the patient first arrives at the hospital. This therapy usually consists of a broad-spectrum antibiotic designed to treat a variety of infections, since the infection-causing organism has not been speciated. This can have unfortunate consequences, such as killing beneficial gut microbiota. Definitive antimicrobial therapy is when certain antibiotics are chosen after culturing microorganisms obtained from the infection site and determining the causative agent of disease. These antibiotics are typically more narrow spectrum in nature and are designed to treat specific infections, but unfortunately, are not able to be given to the patient until the diagnosis is confirmed, leading to the loss of valuable time needed when combating infections effectively (Leekha *et al*. 2011). This delay in prescription of narrow spectrum antibiotics is due to the acknowledgment that early prescription without confirmation of diagnosis can result in treatment failure. Clinicians are also discouraged from prescribing narrow spectrum antibiotics without diagnosis confirmation due to antimicrobial stewardship efforts to prevent over prescription leading to increased antibiotic resistance. There are current recommendations for antibiotic combination therapies for some species, such as vancomycin with an aminoglycoside and rifampin for methicillin-resistant *Staphylococcus aureus* (Hassoun *et al*. 2017), and ceftazidime and gentamicin (sometimes in combination with rifampin) for infections caused by *Pseudomonas aeruginosa* (Ghani *et al*. 1997, Morgan 2014). Combinations of antibiotics are chosen based on the assumption that synergism is conserved between species and strains of microorganisms. However, recent studies have shown that these interactions can change depending on both the species present and the antibiotics prescribed (Fatsis-Kavalopoulos *et al*. 2020). For example, antibiotic combination of ciprofloxacin with metronidazole has been shown to allow for ciprofloxacin recalcitrance to develop in *P. aeruginosa* as the SOS response is induced (Hocquet *et al*. 2014). Antibiotic tolerance can also be increased when certain species are present, such as when *P. aeruginosa* produces pyocyanin allowing for increased antimicrobial tolerance for both itself and other species (Zhu *et al*. 2019). The microbial community plays a role in determining a species’ antimicrobial susceptibilities. Therefore, it is crucial to consider the community when prescribing antibiotics to treat infections effectively.

The checkerboard assay is one way of measuring the effects of antibiotic combination therapy. Varying concentrations of two antibiotics can be dispensed along the columns and rows to allow for determination of minimum inhibitory concentration (MIC) for each antibiotic in combination (Bellio *et al*. 2021). The MICs are determined using visible turbidity readings or optical density. Unfortunately, using the standard readout of visible turbidity and optical density readings from current checkerboard methods does not include the determination of viable cell numbers. While visible turbidity can demonstrate how a community affects the overall susceptibility of all species combined, it leaves the researcher unable to determine where the susceptibilities lie for individual species within the community. Another limitation of the visible turbidity assay is the sensitivity issue. First, dead cells can falsely contribute to turbidity. Second, live cell populations can be too dilute to be read via turbidity, masking the bacteriostatic versus bactericidal effects of certain drug treatments. The new methodology described herein allows for the determination of viable cell numbers via the use of colony forming unit (CFU) counts as the readout for the checkerboard assay, thus allowing for measurement of susceptibility changes for individual species in a polymicrobial community. No special laboratory equipment is needed, only selective/differential media for the species tested. The addition of accurate CFU counts to the data obtained from the checkerboard assay is essential to helping understand exactly how antibiotic combination affects clearing of pathogens known to cause infection when found in a polymicrobial community. As shown with our data, when CFU counts are obtained, even recommended antimicrobial combinations that work well on pathogens by themselves, such as gentamicin with ceftazidime for *P. aeruginosa*, can be antagonistic and counterproductive to patient treatment when a complex microbial community is present.

## MATERIALS AND METHODS

### Bacterial Isolates and Media

The four isolates used in these experiments were *S. aureus* ATCC^®^ 29213, *P. aeruginosa* ATCC^®^ 27853, *E. faecalis* ATCC^®^ 29212, and *A. baumannii* ATCC^®^ 19606. *S. aureus* grew yellow colonies on Mannitol Salt Agar (Fisher Scientific™). *P. aeruginosa* grew green colonies on *Pseudomonas* Isolation Agar (Fisher Scientific™). *E. faecalis* grew black colonies on Bile Esculin Agar with Azide (VWR^®^). *A. baumannii* grew pink colonies on Leeds (VWR^®^).

### Preparation of microbial overnight cultures

A sterile loop was used to obtain approximately five colonies from a selective/differential plate. The colonies were then inoculated into a 125 ml glass flask containing 5 ml of LB and incubated for 18 hours in a shaking incubator at 200 rpm and 37°C under ambient air conditions. After 18 hours of incubation, the overnight cultures contained approximately 10^9^ CFU ml ^-1^ as shown by diluting and plating on the appropriate selective/differential media for each species.

### Antibiotic Preparation and Dilution

All wells in a 96-well plate were filled with 100 μl of Cation-Adjusted Mueller Hinton broth (CAMHB). A 256 μg ml^-1^ storage stock was created for each antibiotic, using methods from the Clinical Laboratory and Standards Institute (CLSI) manual (CLSI M7 2018, CLSI M100 2018). 100 μl of 256 μg ml^-1^ concentration antibiotic stock for antibiotic A was added to all of the wells in the first column of the 96-well plate. 100 μl from that first column was then diluted using serial dilutions across the rest of the wells, all the way to column 11 (1:2 dilutions starting with a concentration of 128 μg ml^-1^ and ending with a final concentration of 0.125 μg ml^-1^). This process was then repeated in a separate 96-well plate for antibiotic B (Sup. Fig. 1).

### Preparation of microbial inoculum

A spectrophotometer was used to measure the OD_625_ of each bacterial species as per CLSI protocol (CLSI M7 2018, CLSI M100 2018). The bacteria were then diluted in 1X PBS to a McFarland standard of 0.08. Next, 40 μl of the McFarland standard was added to 760 μl 1X PBS to create the inoculums for each species. For the polymicrobial inoculum, the 40 μl was split evenly between the 4 species (10 μl per species). Inoculum CFU ml^-1^ obtained from plating on selective/differential media were reported (Sup. Tab. 1).

### Checkerboard assay setup

In two empty 96-well plates, 45 μl of each antibiotic was added at one concentration higher than desired, with one antibiotic in columns, and one in rows (for example, when a concentration of 0.25 μg ml^-1^ of gentamicin was desired, 45 μl of 0.5 μg ml^-1^ gentamicin would be added to the well in addition to 45 μl of the other desired concentration of ceftazidime) (Sup. Fig. 2). The 128 μg ml^-1^ wells were filled from the 256 μg ml^-1^ antibiotic stocks. Each species had its own checkerboard, and the community was given its own checkerboard (Sup. Fig. 3). At the edge of one of the panels, five wells were filled for the growth control (one per species and one for the community), five wells for the vehicle control (sodium carbonate plus water for ceftazidime), and two wells for the negative contamination check (one with the vehicle and one without). All wells except for the negative contamination check wells were inoculated with 10 μl of the appropriate microbial inoculums, and the checkerboards were then incubated for 18 hours. At this time, each inoculum was diluted out ranging from 10^-1^ to 10^-8^ in 1X PBS and then 5 μl of each dilution was plated on selective/differential media. The inoculum plates were incubated along with the panels to get CFU ml^-1^ of the inoculum.

### Preparation of 96-well microplates for dilutions

A sterile pipetting reservoir was filled with 1X PBS. For the 1:10 dilution (first dilution), 90 μl of 1X PBS was added to each well of a 96-well plate. For the 1:100 dilution (second dilution), 198 μl of 1X PBS was added to each well in a 96-well plate.

### Diluting and plating the checkerboard

After 18 hours, the checkerboards were read for visible turbidity. All wells were then diluted twice by pipetting 10 μl into 90 μl (1:10 dilution), followed by 2 μl into 198 μl (1:100 dilution) for a total dilution factor of 1:1000. Then, 5 μl of each of these dilutions were plated on selective and differential media (Sup. Fig. 4). After incubation for 18-24 hours, colony counts were obtained for each species in both the polymicrobial and monomicrobial conditions at each concentration for each antibiotic represented in the checkerboards. CFU counts were multiplied by two to be representative of plating 10 μl.

### Data analysis

All doubled CFU counts were recorded in an Excel spreadsheet representing the checkerboard wells. Biological repeats were averaged together to get the average CFU for each well (180 CFU was substituted for wells that were “TNTC” (“too numerous to count”) as an upper limit on CFU counts to enable data averages to be determined among replicates). Conditional formatting was used in Excel to display the varying CFU counts for each well (the higher the CFU count, the darker the color of the well). All data present in the graphs is representative of the average of three replicates. A Two-sample T-test was performed to compare both monomicrobial and polymicrobial data to their respective growth controls for each species, as well as to compare monomicrobial verses polymicrobial CFU counts. P-values < 0.05 were reported as significant.

## RESULTS

When turbidity data was compared to the CFU data in the classic monomicrobial checkerboard setup, it became evident that viable cells could be detected in wells not displaying visible turbidity for all individual species tested (Fig. 1–4). Conversely, the presence of visible turbidity could also obscure the fact that the cells in particular wells are no longer viable. For example, *S. aureus* had detectable CFUs in 64 μg ml^-1^ ceftazidime even though visible turbidity did not extend past 32 μg ml^-1^, whereas these same data showed that the presence of cells could sometimes be visibly detected in 4 μg ml^-1^ gentamicin, but these cells were no longer viable when plated (Fig. 1). *E. faecalis* and *A. baumannii* also exhibited quantifiable viable cell counts where visible turbidity is lacking, although for the most part the turbidity and CFU data were consistent with each other (Fig. 2–3). Even more strikingly, *P. aeruginosa* had a visible turbidity MIC of 1 μg ml^-1^ for gentamicin, yet there were still large quantities of growth in wells containing a concentration of 1-4 μg ml^-1^ (Fig. 4). Even at 8 μg ml^-1^ gentamicin, there was still detectable growth in one of the replicates when CFUs were measured (Fig. 4B). In total, the monomicrobial checkerboard data revealed that CFU data is at least as reliable as the turbidity data. Additionally, CFU data enabled a differentiation between bactericidal and bacteriostatic antibiotic concentrations. Because the initial inoculum was ~3×10^6^ CFU ml^-1^ (Sup. Table 1), the plating strategy outlined in the methods would yield a CFU count of ~3 colonies per well for the initial inoculum. Therefore, antibiotic concentrations resulting in no detectable growth would be considered bactericidal, conditions resulting in low number of colonies (~1-10 CFU) could be considered bacteriostatic since they prevented further growth but did not kill the inoculum, and conditions resulting in numbers greater than 10-20 CFU should be assumed to support bacterial growth even in cases in which growth did not reach a visually detectable turbidity.

**Figure 1.**
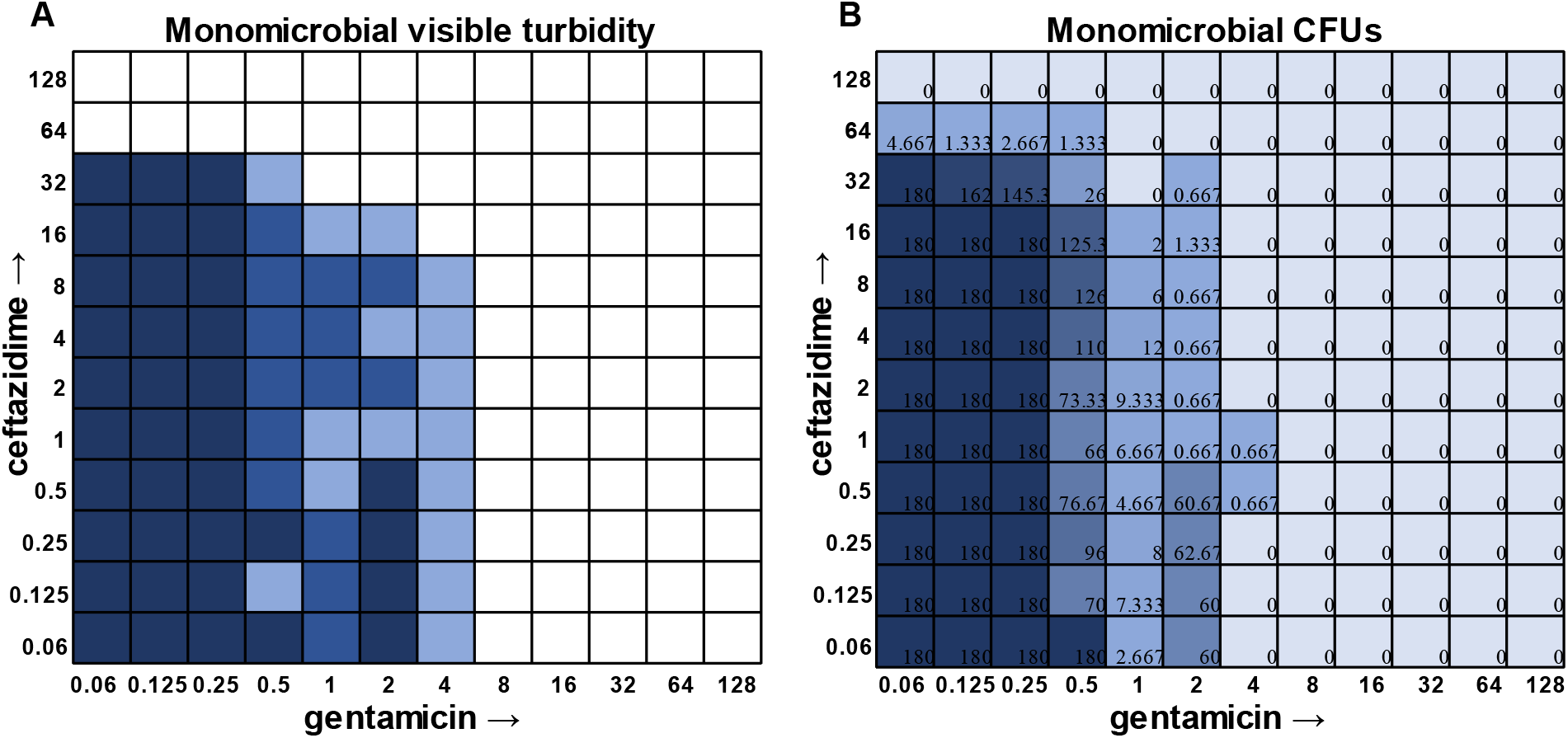
Turbidity versus CFU data in *Staphylococcus aureus* checkerboard assay. **A.** Visible turbidity of *S. aureus* ATCC 29213 when grown in the monomicrobial condition. Data represents triplicate checkerboard assays with ceftazidime and gentamicin. Dark coloration indicates that turbidity was visible in the well in all replicates, medium coloration indicates that turbidity was visible in two replicates, light coloration indicates that turbidity was visible in one replicate. **B.** CFU data from the same triplicate plates with numbers representing the average of the triplicate CFUs detected. Wells displaying TNTC (too numerous to count) CFUs were assigned a number of 180 to enable the determination of averages across replicates.

**Figure 2.**
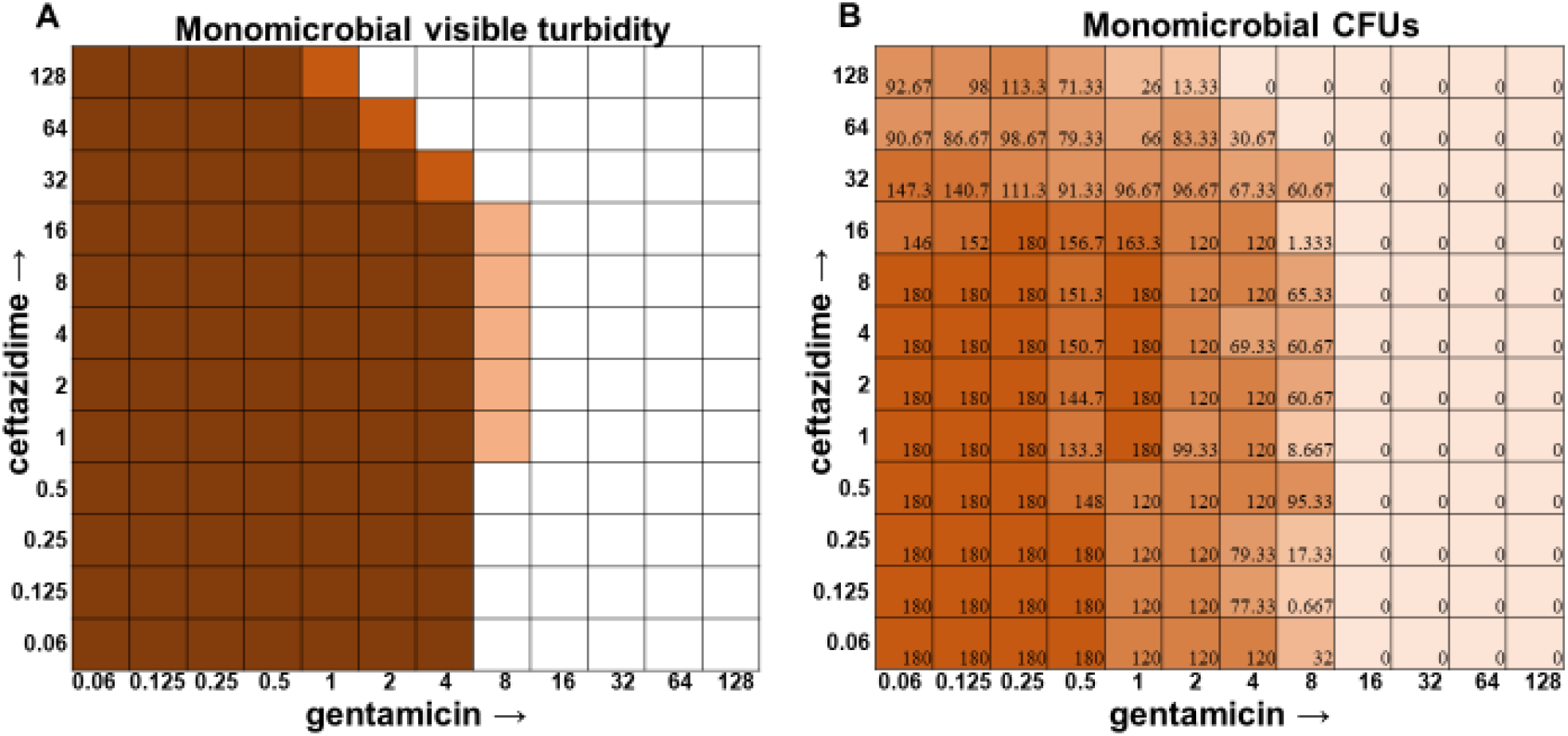
Turbidity versus CFU data in *Enterococcus faecalis* checkerboard assay. **A.** Visible turbidity of *Enterococcus faecalis* ATCC 29212 when grown in the monomicrobial condition. Data represents triplicate checkerboard assays with ceftazidime and gentamicin. Dark coloration indicates that turbidity was visible in the well in all replicates, medium coloration indicates that turbidity was visible in two replicates, light coloration indicates that turbidity was visible in one replicate. **B.** CFU data from the same triplicate plates with numbers representing the average of the triplicate CFUs detected. Wells displaying TNTC (too numerous to count) CFUs were assigned a number of 180 to enable the determination of averages across replicates.

**Figure 3.**
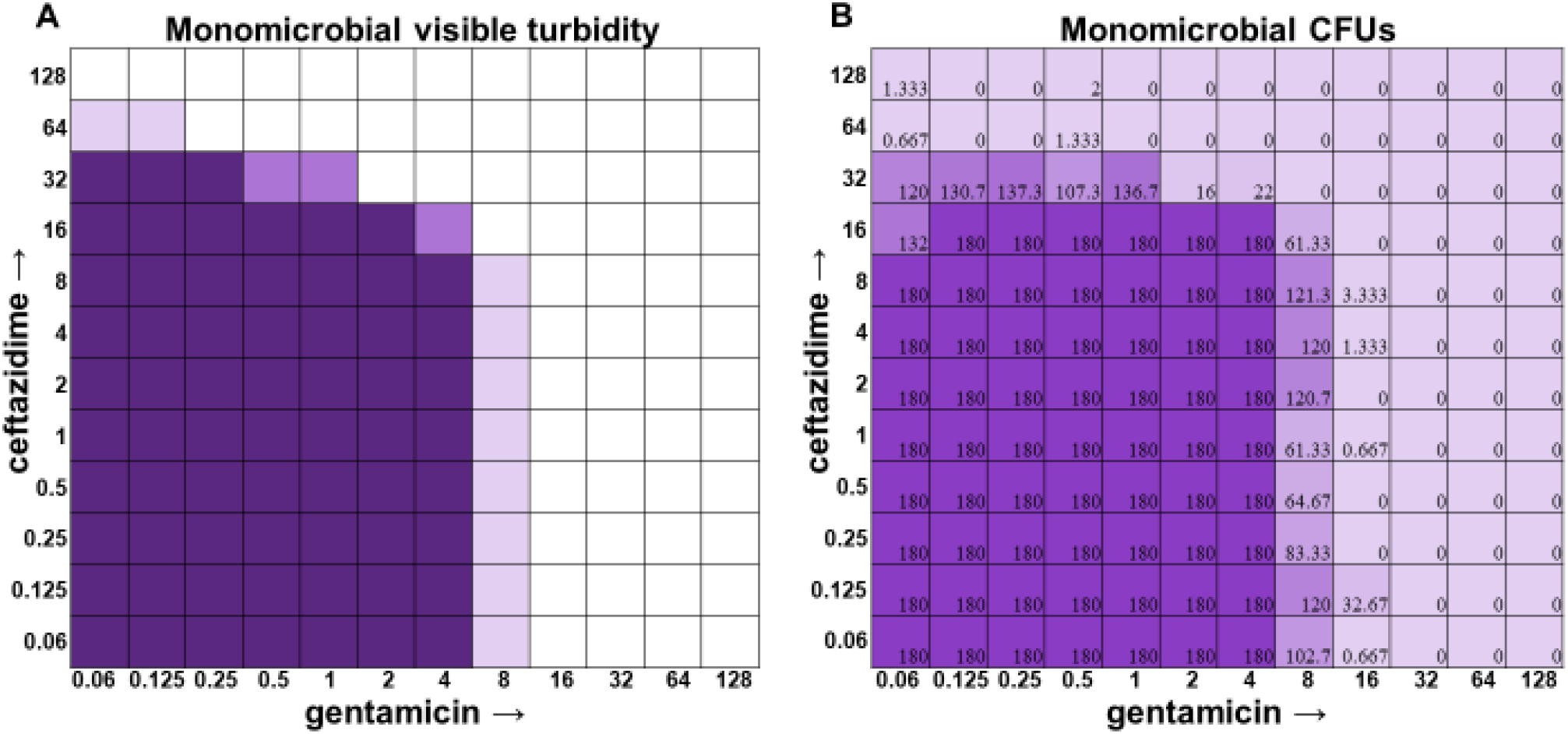
Turbidity versus CFU data in *Acinetobacter baumannii* checkerboard assay. **A.** Visible turbidity of *Acinetobacter baumannii* ATCC 19606 when grown in the monomicrobial condition Data represents triplicate checkerboard assays with ceftazidime and gentamicin. Dark coloration indicates that turbidity was visible in the well in all replicates, medium coloration indicates that turbidity was visible in two replicates, light coloration indicates that turbidity was visible in one replicate. **B.** CFU data from the same triplicate plates with numbers representing the average of the triplicate CFUs detected. Wells displaying TNTC (too numerous to count) CFUs were assigned a number of 180 to enable the determination of averages across replicates.

**Figure 4.**
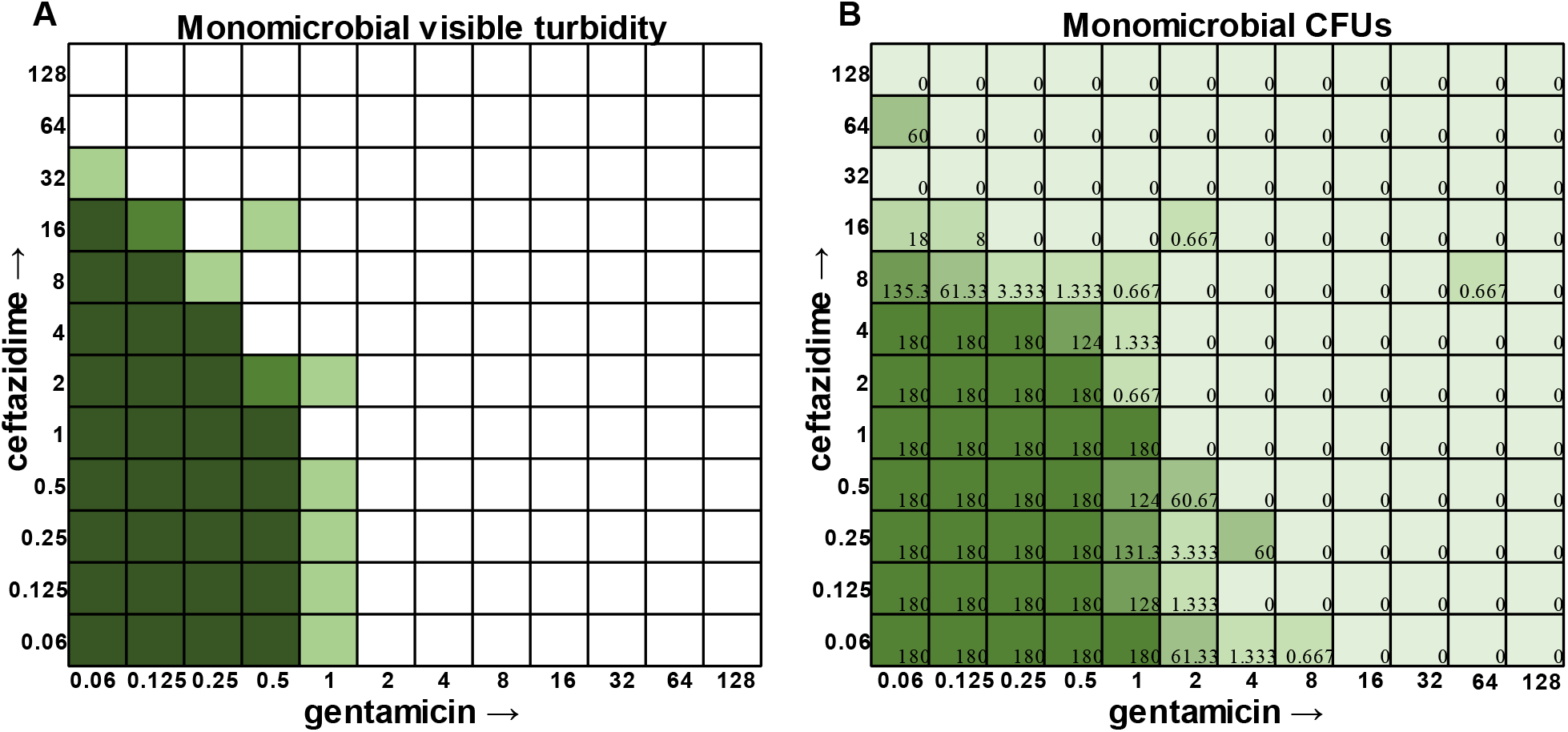
Turbidity versus CFU data in *Pseudomonas aeruginosa* checkerboard assay. **A.** Visible turbidity of *Pseudomonas aeruginosa* ATCC 27853 when grown in the monomicrobial condition. Data represents triplicate checkerboard assays with ceftazidime and gentamicin. Dark coloration indicates that turbidity was visible in the well in all replicates, medium coloration indicates that turbidity was visible in two replicates, light coloration indicates that turbidity was visible in one replicate. **B.** CFU data from the same triplicate plates with numbers representing the average of the triplicate CFUs detected. Wells displaying TNTC (too numerous to count) CFUs were assigned a number of 180 to enable the determination of averages across replicates.

In general, checkerboard assays demonstrate whether two drug interactions will provide a positive combined effect (additive) or a negative combined effect (antagonistic) (Bellio *et al*. 2021). For the purposes of this study, we are defining additive effects as combinations in which the MIC of one antibiotic decreases in the presence of another antibiotic. Antagonistic effects will be defined as combinations in which the MIC of one antibiotic increases in the presence of another antibiotic. Additionally, we will define instances in which the MIC of an antibiotic is unchanged by the presence of another antibiotic as “indifference”. The additive effect of the clinically-used ceftazidime and gentamicin combination (Ghani *et al*. 1997, Morgan *et al*. 2014) is most strikingly observed in *P. aeruginosa* (Fig. 4), which is not surprising considering that this combination is specifically recommended for treatment of *P. aeruginosa* infections. For *S. aureus, E. faecalis*, and *A. baumannii*, the combination of ceftazidime and gentamicin displayed overall MIC indifference with only extremely subtle additive effects (Fig. 1–3).

Once the methodology for cell viability readouts had proved effective in monoculture checkerboard assays, we sought to assess the usefulness of this technique in the polymicrobial community. It should be noted that, in addition to turbidity, we observed that we could visualize *P. aeruginosa* growth via visual detection of its green pigment pyocyanin (Fig. 5 A). However, the cellular source of the turbidity for all other members of the community was indistinguishable, necessitating the use of CFU plating on differential and selective media. Indeed, plating on Mannitol Salt Agar for *S. aureus* (Fig. 5B), Bile Esculin Agar with Azide for *E. faecalis* (Fig. 5C), Leeds for *A. baumannii* (Fig. 5D), and *Pseudomonas* Isolation Agar for *P. aeruginosa* (Fig. 5E) enabled MIC determination for each individual member of the community.

**Figure 5.**
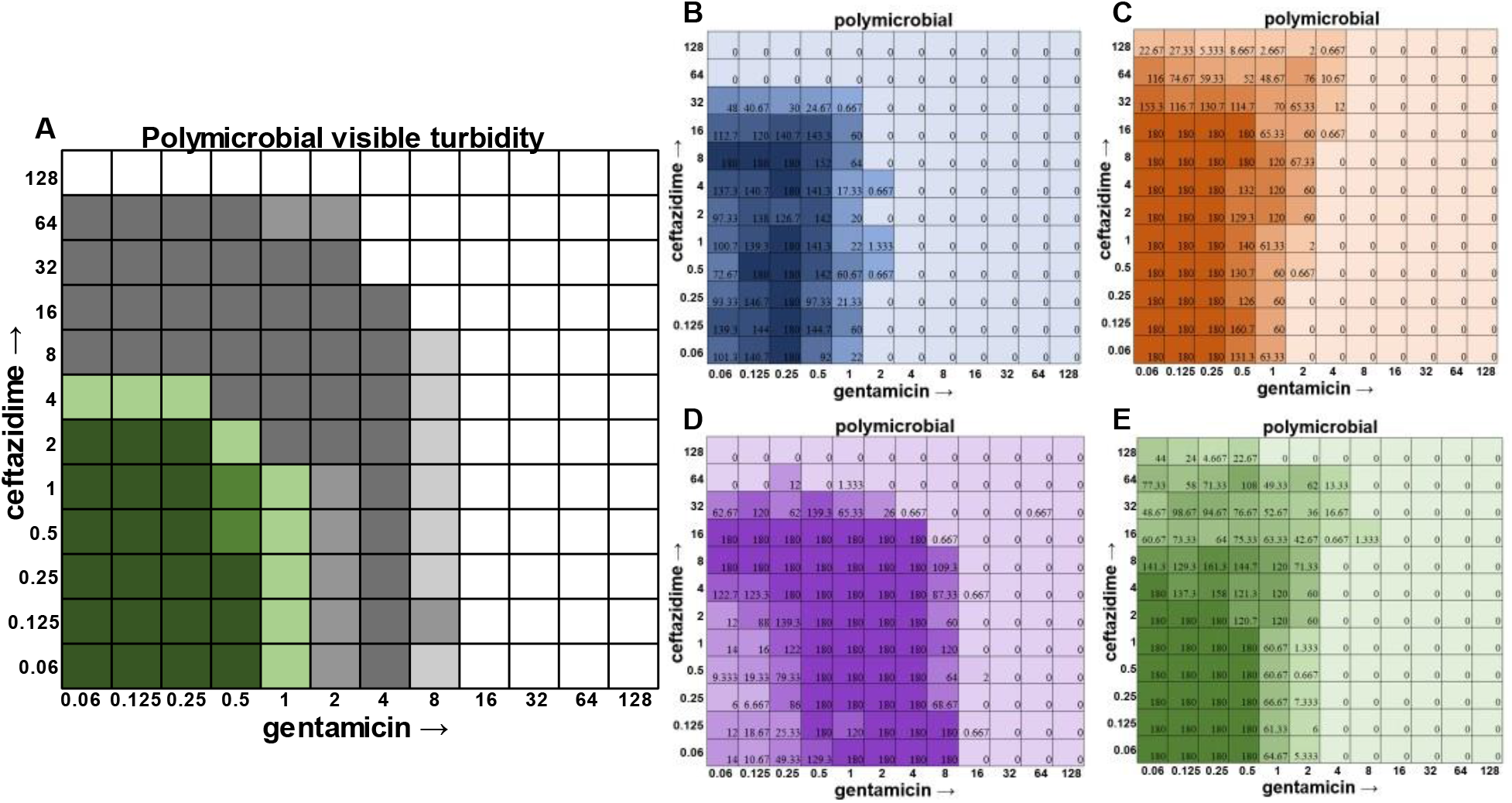
Checkerboard assay with the four species polymicrobial community. **A.** Visible turbidity of the polymicrobial community consisting of all four species. Data represents triplicate checkerboard assays with ceftazidime and gentamicin. Green represents where *P. aeruginosa* pigmentation is present, while gray represents turbidity without green pigmentation. Dark coloration indicates that turbidity was visible in the well in all replicates, medium coloration indicates that turbidity was visible in two replicates, light coloration indicates that turbidity was visible in one replicate. **B-E.** CFU counts from each individual species when grown in the polymicrobial condition are shown on the right (**B.** *S. aureus* in blue, **C.** *E. faecalis* in orange, **D.** *A. baumannii* in purple, and **E.** *P. aeruginosa* in green). As shown above, bacteria are still present even after there is no longer any visible turbidity. Loss of *P. aeruginosa’* green pigmentation is also associated with loss of interspecies competition, as shown by the increase in *A. baumannii’s* population as the pigment disappears from view. Data represent averages of triplicates performed on different days.

In the polymicrobial community, we were able to observe some interspecies competitive behavior. For example, in wells in which *P. aeruginosa* was producing pyocyanin (Fig. 5A), a growth suppression of *A. baumannii* was observed that was reversed when antibiotic concentrations were high enough to inhibit the production of virulence factors in *P. aeruginosa* (Fig 5D). It is worth noting that competition in polymicrobial communities can also make some individuals more susceptible to antibiotics. For example, *S. aureus* became sensitized to gentamicin at 2 μg ml^-1^ and ceftazidime at 64 μg ml^-1^ in the polymicrobial condition relative to monomicrobial culture although overall the combination of the antibiotics exhibited MIC indifference in either culture condition (Fig. 6). Even more strikingly, *E. faecalis* became dramatically sensitized to gentamicin in the polymicrobial context although addition of increasing amounts of ceftazidime antagonized this effect (Fig. 7). Interestingly, the antibiotic MIC for *A. baumannii* appears unaffected by the polymicrobial community even though it was evident that its growth was likely inhibited by its polymicrobial competitor *P. aeruginosa* at lower antibiotic concentrations and the ceftazidime/gentamicin combination for *A. baumannii* displayed relative MIC indifference with only subtle additive effects in either condition (Fig. 8).

**Figure 6.**
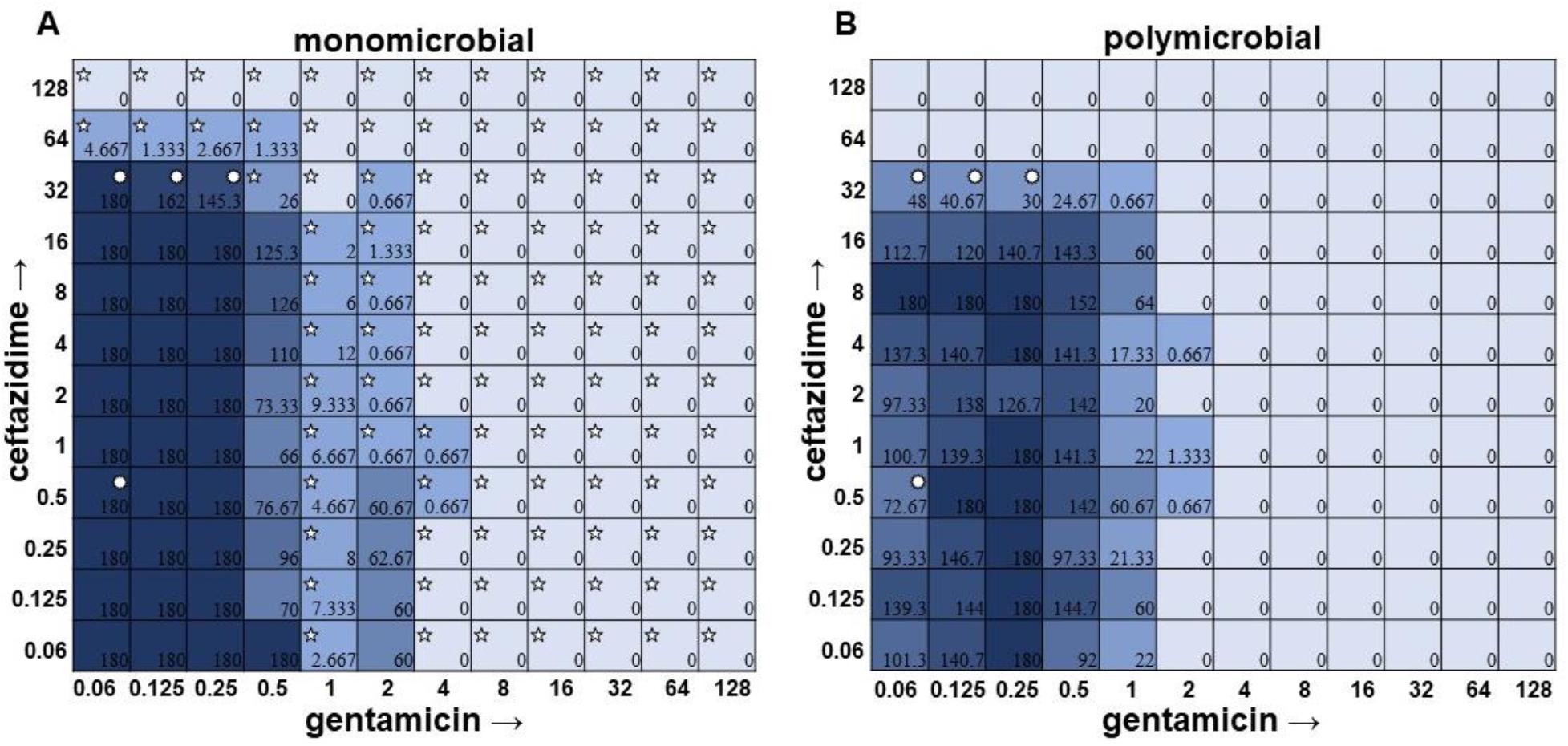
Monomicrobial versus polymicrobial checkerboards reveal that *S. aureus* is slightly sensitized to both ceftazidime and gentamicin in polymicrobial culture. CFU counts for *Staphylococcus aureus* ATCC 29213 when grown in the **A.** monomicrobial condition versus the **B.** polymicrobial condition. As shown above, *S. aureus* becomes sensitized to ceftazidime at 64 μg ml^-1^ and sensitized to gentamicin at 2 μg ml^-1^ in the polymicrobial condition. 5 pointed stars in the top left corner of a well represent a CFU count that is significant as compared to the growth control (180 for monomicrobial and 125.33 for polymicrobial). Circular stars in the top right corner of a well represent a CFU count that is statistically significant between monomicrobial and polymicrobial (two-sample T-test).

**Figure 7.**
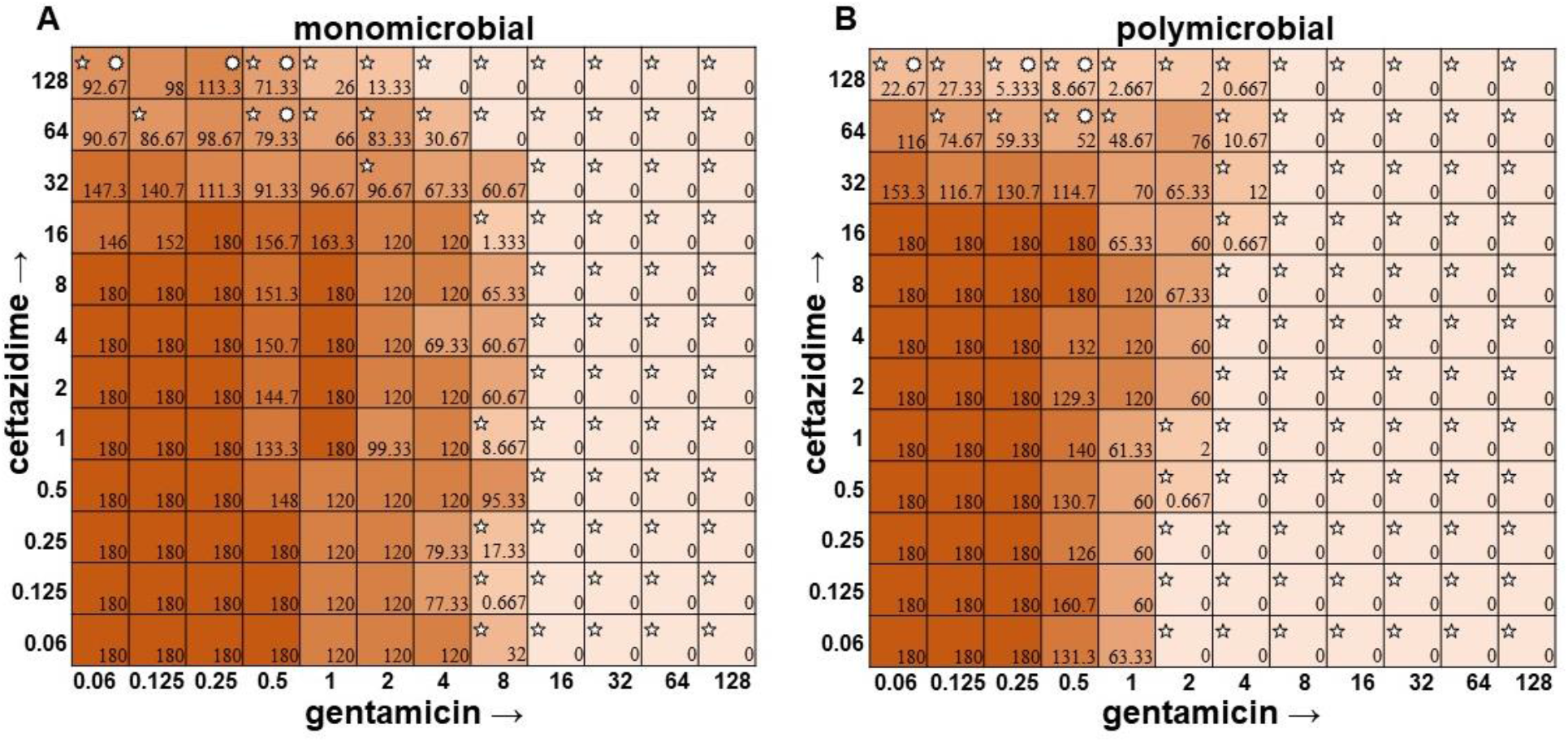
Monomicrobial versus polymicrobial checkerboards reveal that *E. faecalis* is sensitized to gentamicin in polymicrobial culture but this effect is antagonized by ceftazidime. CFU counts for *E. faecalis* ATCC 29212 when grown in the **A.** monomicrobial condition versus the **B.** polymicrobial condition. As shown above, *E. faecalis* becomes sensitized to gentamicin at 2 μg ml ^-1^. 5 pointed stars in the top left corner of a well represent a CFU count that is significant as compared to the growth control (180 for monomicrobial and 180 for polymicrobial). Circular stars in the top right corner of a well represent a CFU count that is statistically significant between monomicrobial and polymicrobial (two-sample T-test).

**Figure 8.**
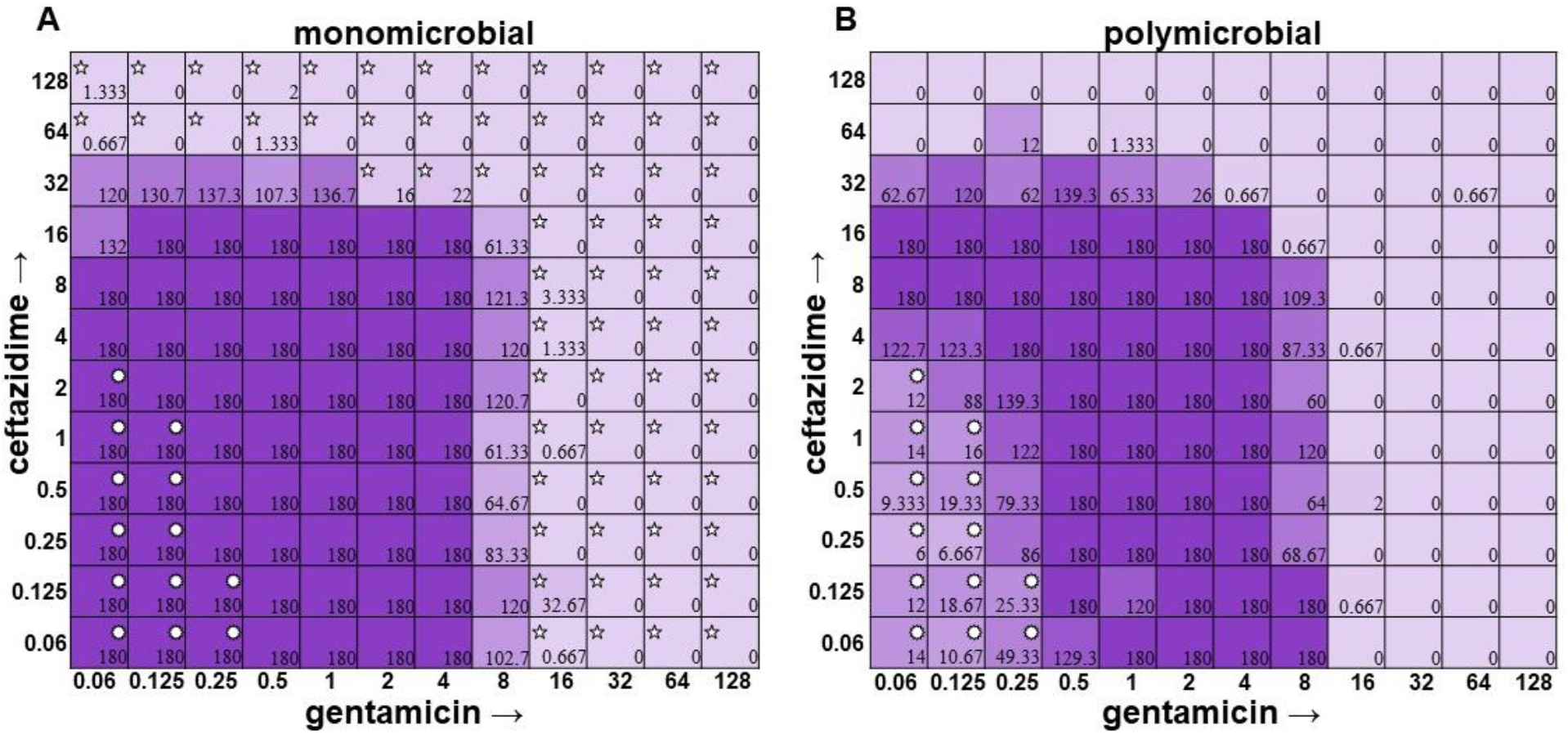
Monomicrobial versus polymicrobial checkerboards reveal that *A. baumannii* succumbs to interspecies competition in low levels of antibiotic but overall MICs are unchanged. CFU counts for *A. baumannii* ATCC 19606 when grown in the **A.** monomicrobial condition versus the **B.** polymicrobial condition. As shown above, *A. baumannii* growth is inhibited in the polymicrobial condition where *P. aeruginosa* growth is present. This is due to antagonization by its competitor. 5 pointed stars in the top left corner of a well represent a CFU count that is significant as compared to the growth control (180 for monomicrobial and 63.33 for polymicrobial). Circular stars in the top right corner of a well represent a CFU count that is statistically significant between monomicrobial and polymicrobial (two-sample T-test).

The most surprising finding of this study was that the gentamicin and ceftazidime combination shown to be effective against *P. aeruginosa* when grown in the monomicrobial condition (Fig. 4; Fig. 9A) became highly antagonistic in polymicrobial culture (Fig. 9B). This is concerning as hospitals base their antimicrobial susceptibility testing on monomicrobial suspensions, which in this case, would be counterproductive to effective patient treatment, as this combination appears to work well when *P. aeruginosa* is grown by itself but is no longer effective when *P. aeruginosa* is present in the community.

**Figure 9.**
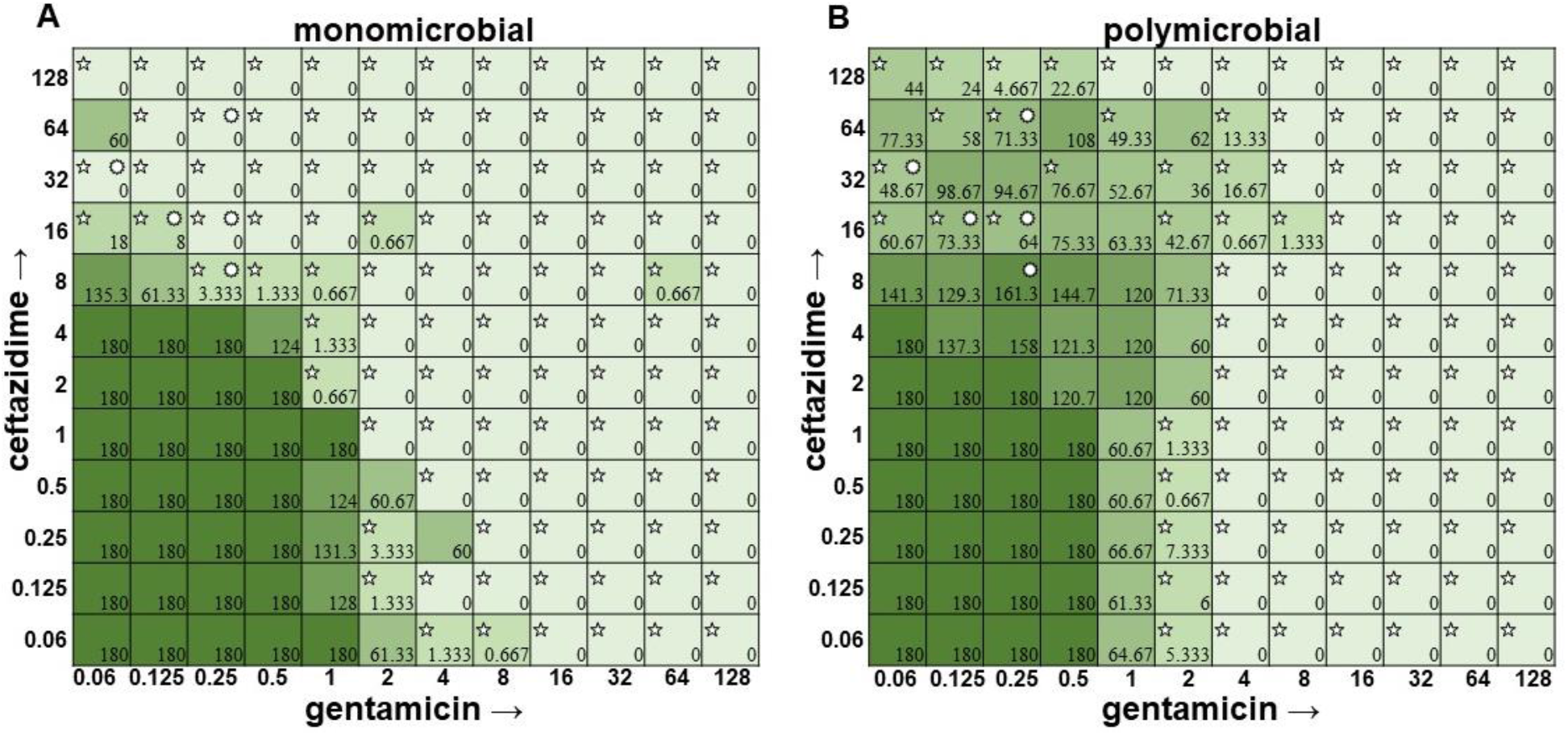
Monomicrobial versus polymicrobial checkerboards reveal that the additive effect of ceftazidime and gentamicin against *P. aeruginosa* becomes antagonistic in polymicrobial culture. CFU counts for *P. aeruginosa* ATCC 27853 when grown in the **A.** monomicrobial condition versus the **B.** polymicrobial condition. As shown above, *P. aeruginosa* becomes desensitized to ceftazidime starting at 8 μg ml ^-1^ and extending up to 128 μg ml ^-1^. This is concerning as ceftazidime is one of the recommended drugs for treating *P. aeruginosa* infections (Babich *et al*. 2020). 5 pointed stars in the top left corner of a well represent a CFU count that is significant as compared to the growth control (180 for monomicrobial and 180 for polymicrobial). Circular stars in the top right corner of a well represent a CFU count that is statistically significant between monomicrobial and polymicrobial (two-sample T-test).

## DISCUSSION

As displayed in the results herein, current checkerboard methodology which focuses on the determination of the MIC of two antibiotics using visible turbidity, cannot account for polymicrobial interactions leading to shifts in the antibiotic susceptibilities of individual species. These shifts can be either antagonistic or additive/beneficial, depending on the antibiotic and the species within the community. This new method does not focus just on visible turbidity, but also accurate CFU counts to be determined for each species in a polymicrobial community, allowing for more accurate changes in individual susceptibilities to be determined. As many infections are polymicrobial in nature (Little *et al*. 2021, Baishya *et al*. 2021), it is important to understand the role the community plays in determining antimicrobial susceptibilities to better treat patient nfections. As the community can be both a help, such as in the case where co-culture with *S. aureus* causes *P. aeruginosa* to alter its biofilm structure to make it resistant to tobramycin (Beaudoin *et al*. 2017), or a hinderance, such as cooperative cross-feeding with anaerobic bacteria causing a now dependent *P. aeruginosa* to become less resilient when challenged with antibiotics (Flynn *et al*. 2020).

Determination of polymicrobial MICs is especially important in cases where the community renders current combinatorial therapies either useless or counterproductive. In the case of gentamicin and ceftazidime, a currently recommended therapy for *P. aeruginosa* (Ghani *et al*. 1997, Morgan *et al*. 2014), this combination becomes antagonistic when *P. aeruginosa* is present in a polymicrobial community. Antimicrobial susceptibility testing (AST) done with the monomicrobial culture of *P. aeruginosa* would reveal an MIC of 32 μg ml^-1^ of ceftazidime needed to completely clear the infection. However, when the bacteria are present in a community, ceftazidime has an MIC >128 μg ml^-1^ against *P. aeruginosa* (Fig. 9). Additionally, if a patient with a polymicrobial infection containing *P. aeruginosa* were to be prescribed a combination of gentamicin and ceftazidime to treat their chronic infection with the dosage based on monomicrobial MICs determined via visible turbidity, our data shows that the addition of higher concentrations of ceftazidime could reverse the usefulness of gentamicin (Fig. 9B). As the effectiveness of treatment influences patient morbidity and mortality, we propose that it is crucial to consider the role of the community a pathogen is in when determining the best treatment. Future directions for this project include the determination of essential community members and the role they play in protecting *P. aeruginosa* from the combinatorial therapy of gentamicin and ceftazidime.

As time is often a concern with treating persistent infections (the longer an infection lasts, the more at risk a patient becomes for complications), it would be undesirable to implement new techniques in the clinical microbiology laboratory that require additional time. Our new method for determining the CFU counts of the individuals in a polymicrobial community focuses on effectively determining the role of the community within the same time frame as one might determine the MIC of a monomicrobial AST panel. As shown in Fig. 10, instead of overnight culturing to determine the causative agent of infection in pure culture microbiology, patient samples could be placed directly into an AST panel and plated after 18 hours on selective/differential media to visualize the different members of the community contributing to changes in antimicrobial susceptibilities as well as the one causative agent of infection. This new methodology would not significantly increase the time needed for MIC determination and allows for better analysis of how the community affects antibiotic susceptibilities, therefore leading to better determination of MICs and more effective prescription of treatment.

**Figure 10.**
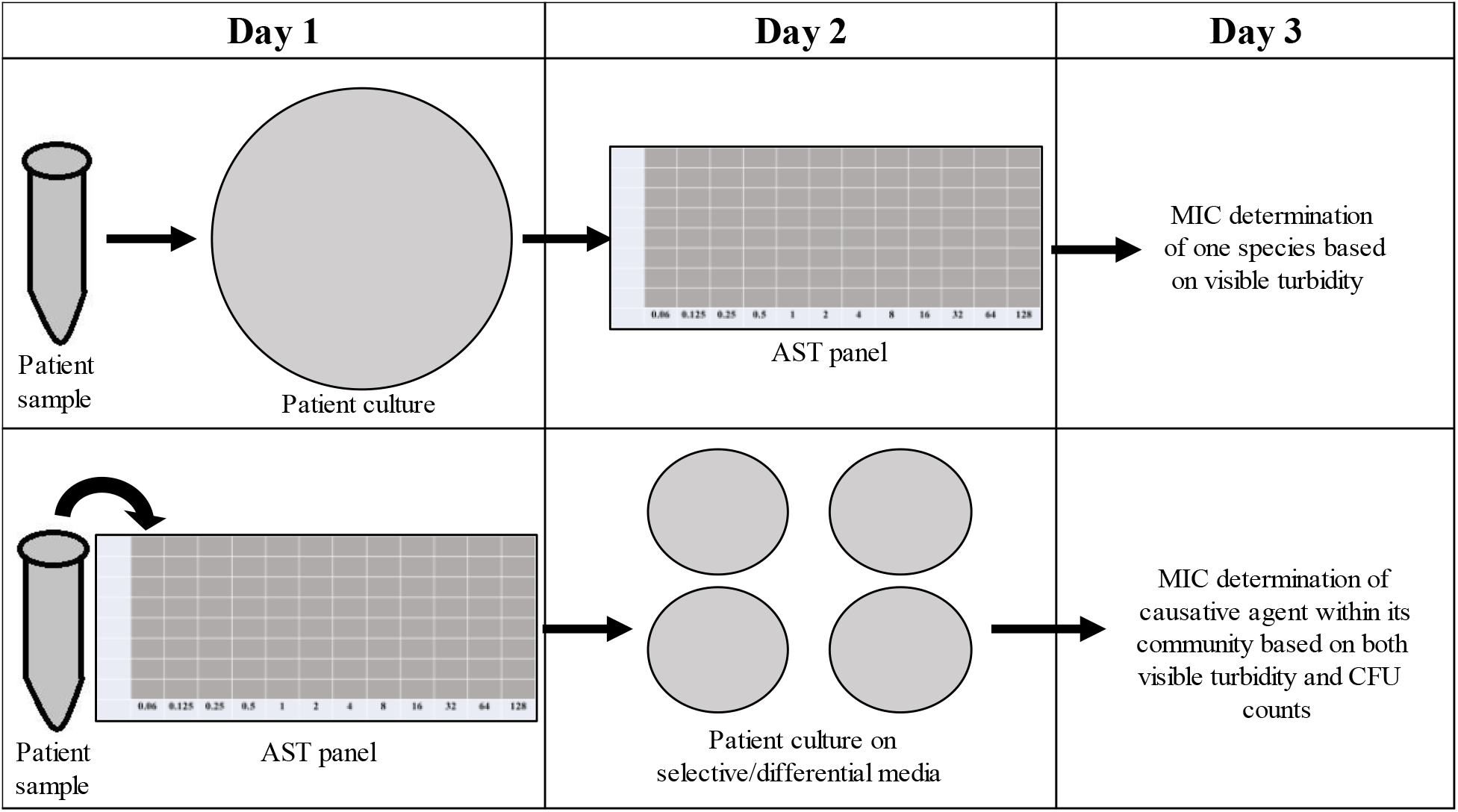
New checkerboard methodology allows for determination of CFUs in a polymicrobial community without extending time needed to determine antibiotic susceptibilities. This represents a comparison of the current method of antimicrobial susceptibility testing (AST) (top row) verses our proposed AST methods (bottom row). As shown in the figure, our method does not extend testing time, yet allows for a more accurate determination of the MIC of antimicrobials used to treat the causative agent. This method not only accounts for the role of the community in determining MIC, but also allows for CFU counts to be obtained, providing an accurate depiction of growth even when visible turbidity shows none.

---

**Supplemental Figure 1.**
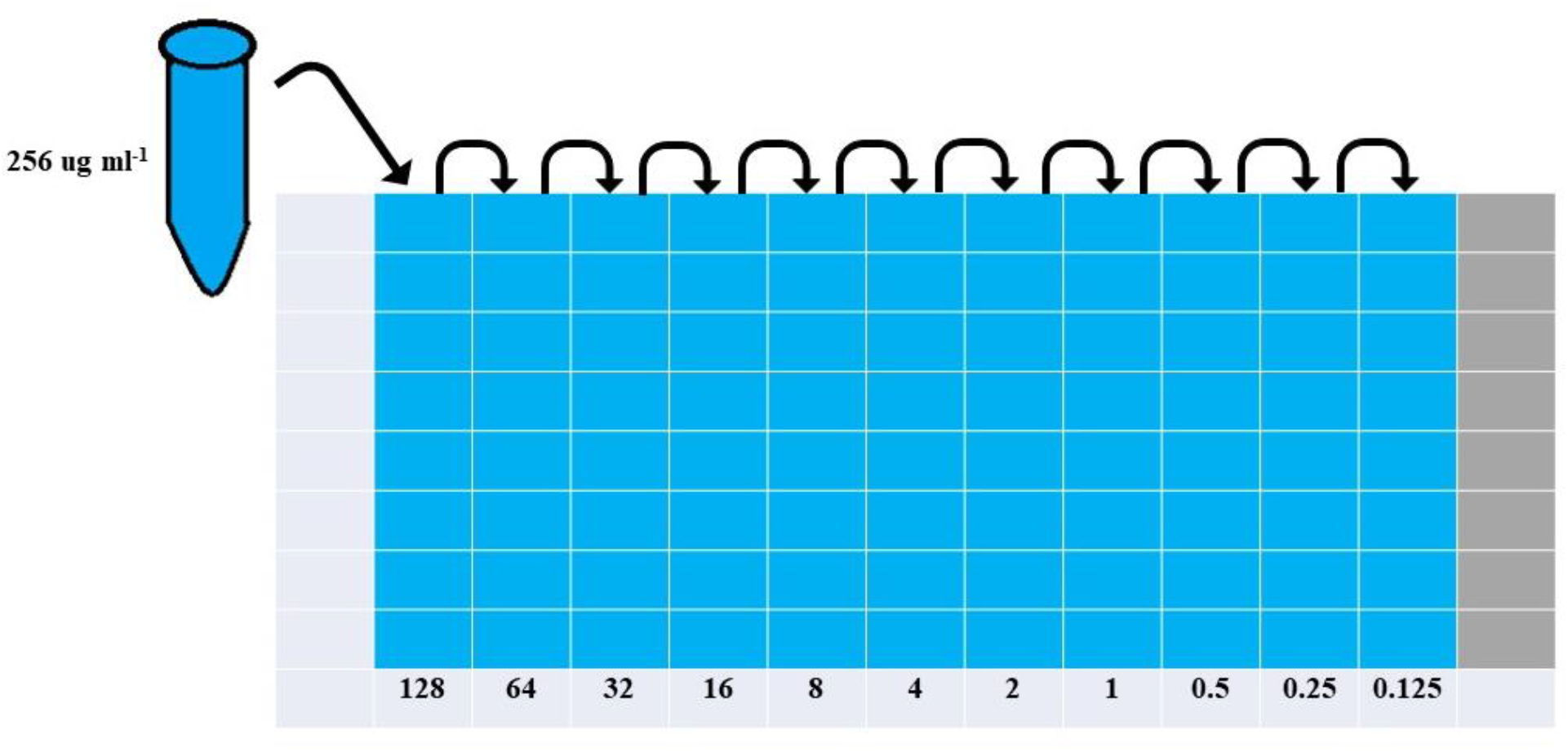
Methods for diluting both antimicrobials start with a 256 μg ml ^-1^ stock, followed by 1: 2 dilutions to a concentration of 0.1 2 5 μg ml ^-1^. Depiction of serial dilutions performed for both antimicrobials. 100 μl of a 256 μg ml ^-1^ stock was diluted out in 100 μl of CAMHB (1:2 dilutions) to a final concentration of 0.125 μg ml ^-1^. 45 μl of each dilution was then added to the checkerboard itself.

**Supplemental Figure 2.**
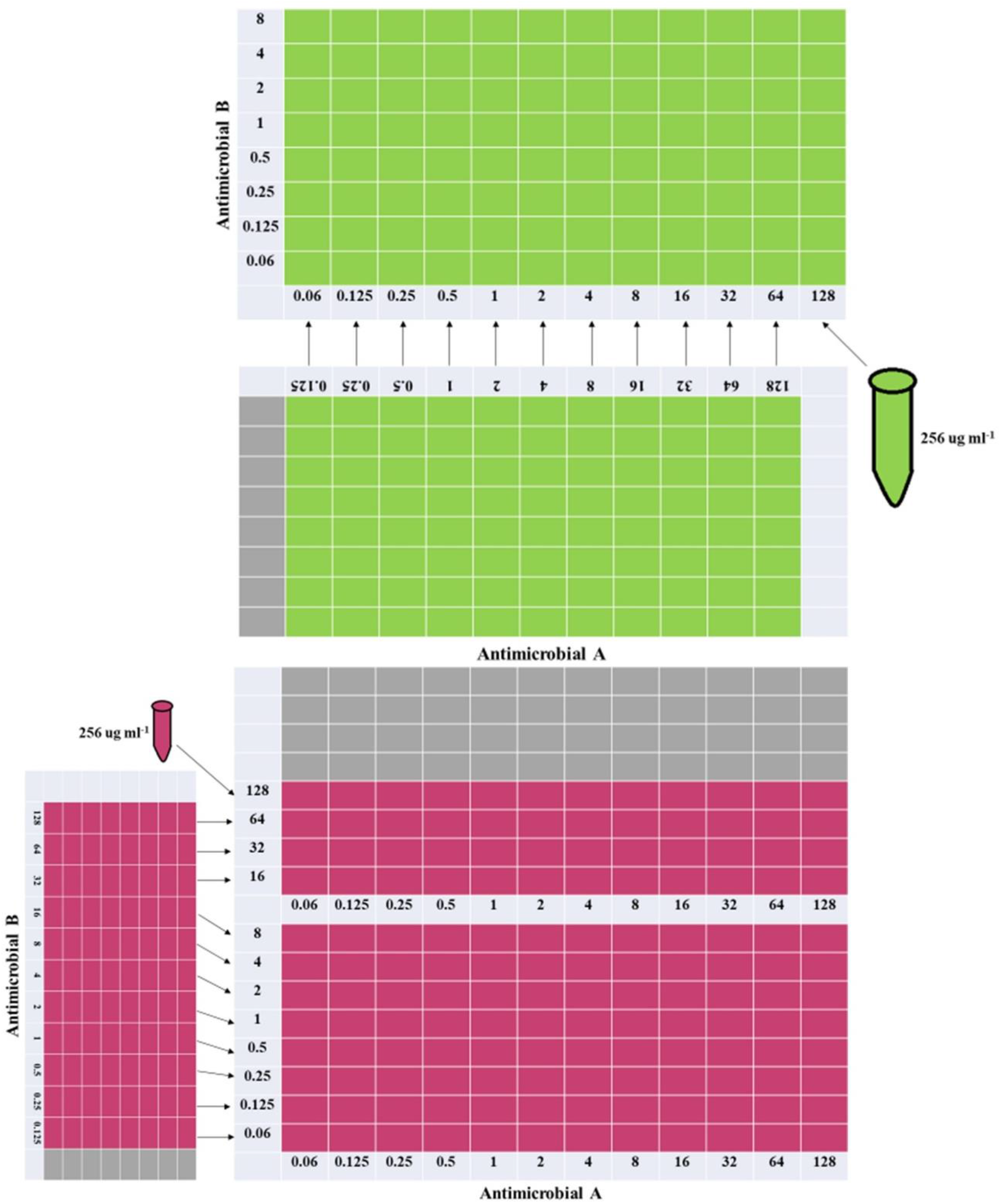
Checkerboard setup has one antimicrobial in rows and one antimicrobial in columns, both with varying concentrations. Each well in the checkerboard received 45 μl of each antimicrobial from a well one concentration higher than desired. For example, if a concentration of 4 μg ml ^-1^ was desired, 45 μl would be taken from the 8 μg ml ^-1^ well.

**Supplemental Figure 3.**
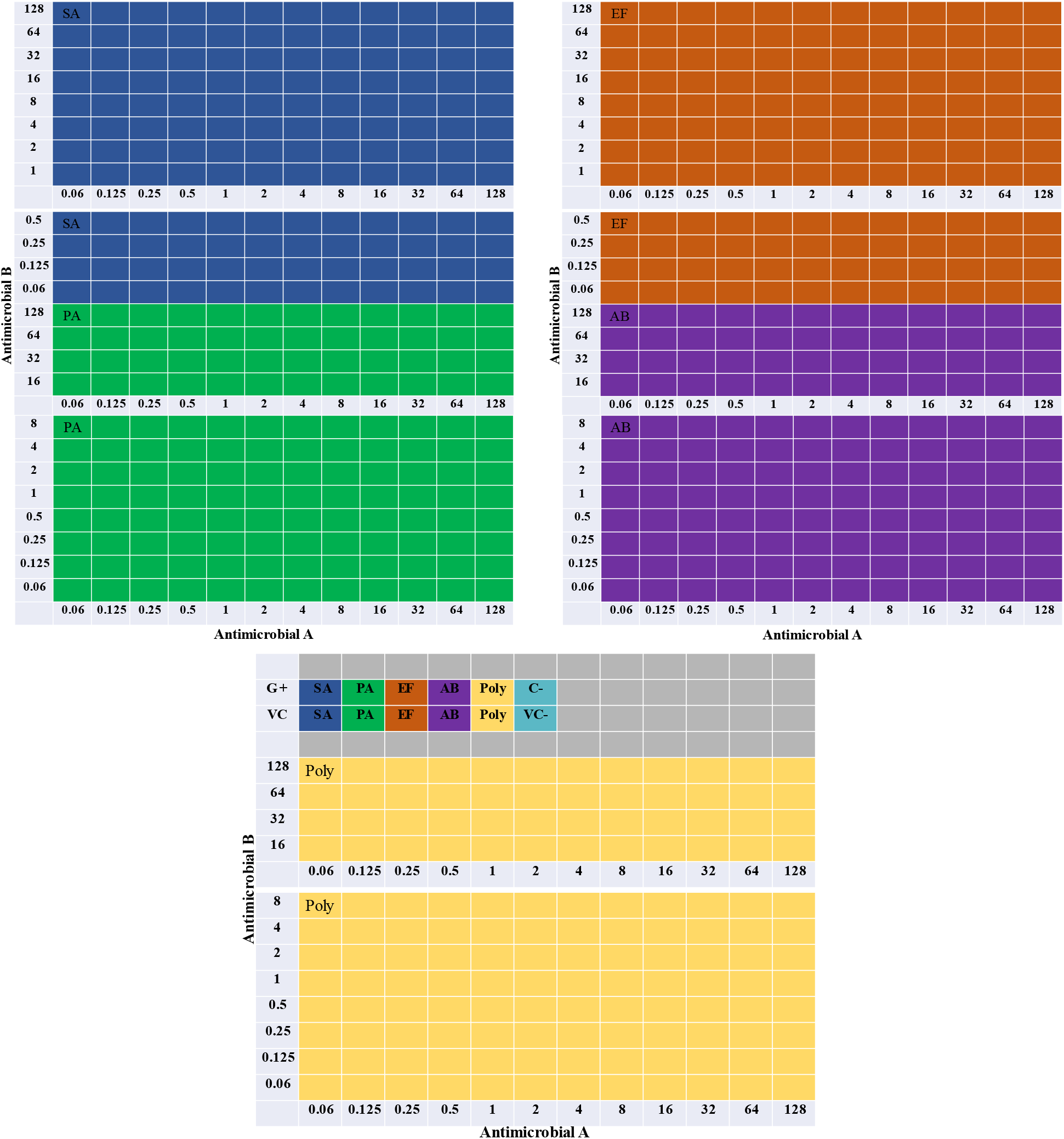
The completed checkerboard setup for these experiments consisted of eight 96-well plates with the four species and the community split between them. Depiction of the complete checkerboard shows each species had 144 wells of antibiotic split across two 96-well plates, a growth control well, and a vehicle control well. A negative contamination check with and without the vehicle was also present on another 96-well plate.

**Supplemental Figure 4.**
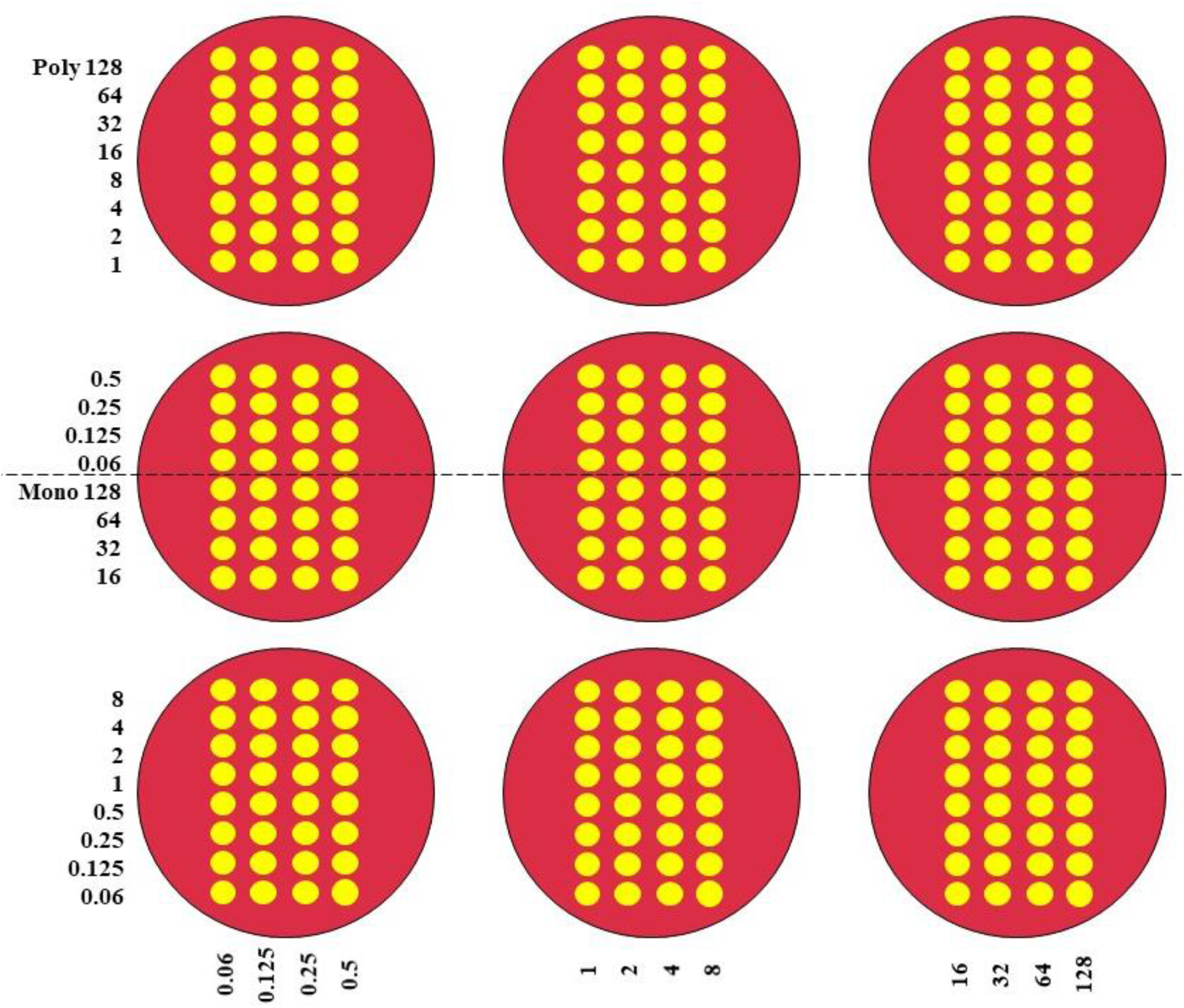
CFUs were obtained by plating 5 μl onto selective/differential media, and then incubating for 18-24 hours. Depiction of plate setup. 5 μl plated onto selective/differential media. The above setup existed for each individual species. The growth control and vehicle control were plated at the edge of a plate. CFU counts were obtained after incubation by multiplying by two to represent what would have been obtained if 10 μl had been plated.

**Supplemental Table 1.** Inoculum CFU ml ^-1^ obtained for each species shows inoculums were very similar across species in the polymicrobial condition. Average CFU ml ^-1^ for each species’ inoculum for both the monomicrobial and polymicrobial conditions. The standard deviation is also included.

| Species | Mono Avg CFU ml<sup>-1</sup> | Mono StDev | Poly Avg CFU ml<sup>-1</sup> | Poly StDev |
| --- | --- | --- | --- | --- |
| SA 29213 | 4.28E+06 | 8.35E+05 | 1.38E+06 | 2.71E+05 |
| PA 27853 | 8.44E+06 | 4.10E+06 | 1.80E+06 | 4.24E+05 |
| EF 29212 | 6.22E+05 | 4.85E+05 | 8.67E+05 | 1.63E+05 |
| AB 19606 | 4.97E+06 | 7.52E+05 | 2.07E+06 | 9.49E+05 |

## REFERENCES

Ahmed, A., Azim, A., Gurjar, M., and Baronia, A. K. (2014). Current concepts in combination antibiotic therapy for critically ill patients. Indian journal of critical care medicine: peer-reviewed, official publication of Indian Society of Critical Care Medicine, 18(5), 310–314.

Angst, D. C., Tepekule, B., Sun, L., Bogos, B., and Bonhoeffer, S. (2021). Comparing treatment strategies to reduce antibiotic resistance in an in vitro epidemiological setting. Proceedings of the National Academy of Sciences, 118(13), e2023467118.

Babich, T., Naucler, P., Valik, J. K., Giske, C. G., Benito, N., Cardona, R., … Yahav, D. (2020). Ceftazidime, Carbapenems, or Piperacillin-tazobactam as Single Definitive Therapy for Pseudomonas aeruginosa Bloodstream Infection: A Multisite Retrospective Study. Clinical Infectious Diseases, 70(11), 2270–2280.

Baishya, J., Bisht, K., Rimbey, J. N., Yihunie, K. D., Islam, S., Al Mahmud, H., Waller, J. E., and Wakeman, C. A.. (2021). The Impact of Intraspecies and Interspecies Bacterial Interactions on Disease Outcome. Pathogens, 10(2), 96.

Beaudoin, T., Yau, Y., Stapleton, P. J., Gong, Y., Wang, P. W., Guttman, D. S., & Waters, V. (2017). Staphylococcus aureus interaction with Pseudomonas aeruginosa biofilm enhances tobramycin resistance. NPJ biofilms and microbiomes, 3, 25.

Bellio, P., Fagnani, L., Nazzicone, L., and Celenza, G. (2021). New and simplified method for drug combination studies by checkerboard assay. MethodsX, 8, 101543.

CLSI M7. Methods for Dilution Antimicrobial Susceptibility Tests for Bacteria That Grow Aerobically. 11th ed. CLSI standard M07. Wayne, PA: Clinical and Laboratory Standards Institute; 2018.

CLSI M100. Performance Standards for Antimicrobial Susceptibility Testing. 28th ed. CLSI supplement M100. Wayne, PA: Clinical and Laboratory Standards Institute; 2018.

Fatsis-Kavalopoulos, N., Roemhild, R., Tang, P.-C., Kreuger, J., and Andersson, D. I. (2020). CombiANT: Antibiotic interaction testing made easy. PLOS Biology, 18(9), e3000856.

Flynn, J. M., Cameron, L. C., Wiggen, T. D., Dunitz, J. M., Harcombe, W. R., & Hunter, R. C. (2020). Disruption of Cross-Feeding Inhibits Pathogen Growth in the Sputa of Patients with Cystic Fibrosis. mSphere, 5(2), e00343–20.

Ghani, M., and Soothill, J. S. (1997). Ceftazidime, gentamicin, and rifampicin, in combination, kill biofilms of mucoid Pseudomonas aeruginosa. Canadian Journal of Microbiology, 43(11), 999–1004.

Hassoun, A., Linden, P. K., and Friedman, B. (2017). Incidence, prevalence, and management of MRSA bacteremia across patient populations—a review of recent developments in MRSA management and treatment. Critical Care, 21(1).

Hocquet, D., and Bertrand, X. (2014). Metronidazole increases the emergence of ciprofloxacin- and amikacin-resistant Pseudomonas aeruginosa by inducing the SOS response. Journal of Antimicrobial Chemotherapy, 69(3), 852–854.

Leekha, S., Terrell, C. L., and Edson, R. S. (2011). General Principles of Antimicrobial Therapy. Mayo Clinic Proceedings, 86(2), 156–167.

Little, W., Black, C., and Smith, A. C.. (2021). Clinical Implications of Polymicrobial Synergism Effects on Antimicrobial Susceptibility. Pathogens, 10(2), 144.

Micek, S. T., Lloyd, A. E., Ritchie, D. J., Reichley, R. M., Fraser, V. J., and Kollef, M. H. (2005). Pseudomonas aeruginosa Bloodstream Infection: Importance of Appropriate Initial Antimicrobial Treatment. Antimicrobial Agents and Chemotherapy, 49(4), 1306–1311.

Morgan, A. E. (2014). The synergistic effect of gentamicin and ceftazidime against Pseudomonas fluorescens. Bioscience Horizons: The International Journal of Student Research, 7.

Zhu, K., Chen, S., Sysoeva, T. A., and You, L. (2019). Universal antibiotic tolerance arising from antibiotic-triggered accumulation of pyocyanin in Pseudomonas aeruginosa. PLOS Biology, 17(12), e3000573.

